# Motor effort and adaptive sampling in perceptual decision-making

**DOI:** 10.1101/637017

**Authors:** Tianyao Zhu

**Affiliations:** Faculty of Education and Psychology, Eötvös Loránd University, Budapest, Hungary

## Abstract

People usually switch their attention between the options when trying to make a decision. In our experiments, we bound motor effort to such switching behavior during a two-alternative perceptual decision-making task and recorded the sampling patterns by computer mouse cursor tracking. We found that the time and motor cost to make the decision positively correlated with the number of switches between the stimuli and increased with the difficulty of the task. Specifically, the first and last sampled items were chosen in an attempt to minimize the overall motor effort during the task and were manipulable by biasing the relevant motor cost. Moreover, we observed the last-sampling bias that the last sampled item was more likely to be chosen by the subjects. We listed all possible Bayesian Network models for different hypotheses regarding the causal relationship behind the last-sampling bias, and only the model assuming bidirectional dependency between attention and decision successfully predicted the empirical results. Meanwhile, denying that the current decision variable can feedback into the attention switching patterns during sampling, the conventional attentional drift-diffusion model (aDDM) was inadequate to explain the size of the last-sampling bias in our experimental conditions. We concluded that the sampling behavior during perceptual decision-making actively adapted to the motor effort in the specific task settings, as well as the temporary decision.

## Introduction

When people try to choose between two similar products in a shopping center, they often approach each shelf where the products are displayed to have a closer look. If the choice is difficult to make, people may walk back-and-forth the two shelves for a long time. Many people will start by examining the product around the entrance of the shop, but choose the one near the checkout counter eventually to save some effort. This daily example suggests that our decisions are not solely shaped by the relative values of the alternatives, but also other factors including the motor effort related to the sampling and the action execution processes.

However, sensorimotor aspects have not been integrated into decision-making studies until recently. It is still an on-going controversy whether action is part of decision-making: According to the Embodied Choice model, action execution is part of the decision-making process rather than merely a means to report the decision; in other words, action can feedback into the decision-making process [1]. Researchers have also studied decision-making by analyzing movement patterns [2] and sought neural imaging evidence for the involvement of the sensorimotor system during decision-making [3].

Meanwhile, Aczel et al. [4] argued that the observed decision bias was not caused by the movement toward one of the options, as the Embodied Choice model proposed, but rather the difference in the required motor effort during action. Other studies also reported the influence of motor effort during action upon decision-making: For example, perceptual decisions have been observed to be biased by the difference in the motor cost to make the response [5]. Moreover, the exposure to the unequal motor cost also biased the subsequent decisions even when they were vocally reported, indicating that motor effort can affect decision-making at a stage earlier than action execution [6]. De Lange and Fritsche [7] suggested that motor cost can influence decision-making similarly to rewards. Besides, motor effort can also affect changes of mind during decision-making [8].

Apart from action, the sampling behavior can be accompanied by motor effort as well, especially when the items to choose from are spatially separated. However, no investigation has focused on the influence of motor effort upon sampling. Although in some paradigms two or more visual stimuli were present, the main form of movement involved during sampling was the saccadic eye movement; unlike limb movements, energy costs are not a significant consideration in the planning for saccades [9].

Another issue following the separation of the options in space is the attention allocation during sampling. Typically, the decision-maker switches the attention (by the behavior of switching the gaze) between the options at least once, sometimes multiple times. What is the relationship between attention and decision-making? Several results showed that manipulation of attention biased the decision [10–14]. Under the assumption that attention can influence value integration during decision-making, Krajbich et al. [15] proposed the attentional drift-diffusion model (aDDM). Unlike the traditional drift-diffusion model where the relevant evidence accumulates at a constant rate (the drift rate) within one decision, the aDDM allows the drift rate to change with attention: the option currently being attended (gazed at) shall receive more evidence. Such a model has successfully explained the gaze patterns and several gaze-related biases in preferential and perceptual decisions performed by human subjects [15–17].

Specifically, the aDDM assumes that attention or gaze switches between the options randomly. In fact, there is rare evidence supporting that temporary choices can influence attention allocation. Shimojo et al. [18] reported the gaze cascade effect that gaze was biased toward the finally chosen item during preferential decision-making, yet Krajbich [19] argued that the phenomenon was readily explained by the aDDM and suggested that gaze or attention has a causal effect on choice, but not vice versa.

Under natural circumstances, humans gather information and sample relevant cues with attention and active sensing behaviors (shift of gaze and assisting limb/body movement) [20]. Sampling behavior itself can be regarded as a low-level decision-making process about what information to acquire, as well as where and when [21]. In the current study, we aim to figure out the factors influencing sampling patterns during a basic perceptual decision-making task, especially how sampling behavior adapts to the expected motor effort given the specific environment of the task. We designed a paradigm in which motor effort was bind to the sampling and action execution processes, and manipulated the expected motor cost to examine corresponding changes in the sampling patterns. Additionally, we tested the causal relationship between the temporary decision and the attention allocation strategy during sampling by analyzing a Bayesian Network model and simulating an aDDM.

## Methods

### Paradigm and stimuli

The paradigm was based on a two-alternative perceptual decision-making task in which subjects were asked to decide which of the two groups of black and white dots contained more white ones. Two imaginary circles (diameter 3.5 cm) were located horizontally apart on the upper half of the screen (20 cm between their centers), each containing 100 dots. The dots were either black or white on a 50% gray background. In each trial, we randomly set the proportions of white dots in each group with the following method: First, we separately drew an average proportion *A* from [0.4, 0.6] and a distinction proportion *D* from [0, 0.3]. The proportions of white dots in the two groups would be *A* ± 0.5*D*. Then, we randomly assigned the two calculated proportions to the left and right group, making sure that in 50% trials there were more white dots in the left group.

To bind motor effort to the sampling process, we applied an artificial rule that the sampling quality is in proportion to the distance between the agent and the stimulus. In natural circumstances, it is interpreted as ‘the closer one gets to look at an object, the more details will be seen’, and ‘getting closer’ needs motor effort. In our paradigm, the position and color of each dot were fixed within the trial, but in each frame (frame rate 60 fps) a different set of randomly selected dots were made invisible so that the dots were ‘blinking’ with varying phase and rate. The number of invisible dots in each frame was in proportion to the current distance between the mouse cursor and each dot stimulus, thus the closer the cursor was to the stimulus, the more dots were visible in a certain period (Fig 1B). When the cursor was moved to the leftmost, the left group of dots would become completely visible and static, while the right group would be completely invisible. Therefore, to get better sampling quality, subjects must make some motor effort to move the cursor closer to the stimulus they want to examine.

**Fig 1.**
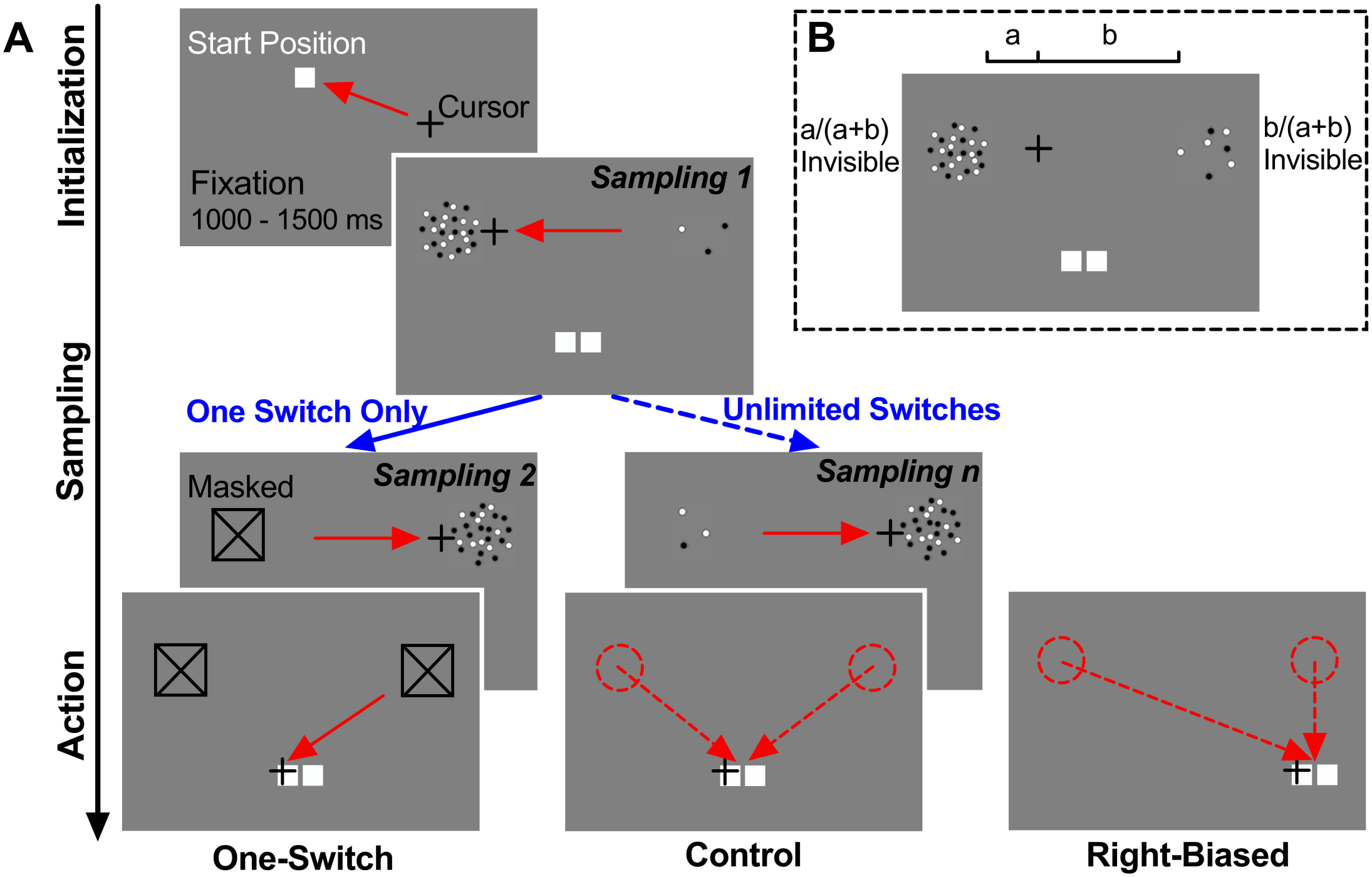
Illustration of the paradigm. **(A)** A fixation (1000 – 1500 ms) on the start position with the mouse cursor was necessary to trigger each trial. After that, subjects moved the cursor to the stimuli alternatively to sample them. Finally, subjects clicked on the corresponding button to report which stimulus contained more white dots. **(B)** The number of invisible dots per frame was in proportion to the distance between the cursor and the stimulus. Subjects must move the cursor close to the stimulus to get better sampling quality.

At the beginning of each trial, a start position was randomly drawn within the central 80% range between the boundaries of the two stimuli, marked by a small white square on the screen. Subjects should drag the computer mouse cursor onto the square and stay fixed for a short time (1000 – 1500 ms randomly) to trigger the trial. After the fixation period, the two stimuli would appear, and the subject could start to sample them. Subjects were told to avoid pausing the cursor in the middle of the screen while looking sideways at the stimuli. We set two sampling modes: In the one-switch mode, subjects should and could only make one switching movement between the stimuli, which means they had only one chance to sample each of the alternatives. When the cursor was moved close enough to the stimulus (visible dots more than 90% per frame) and then left, that stimulus would be masked and could not be examined again in the current trial. Subjects were instructed not to move their mouse to an already masked stimulus. The length of time to examine each stimulus was not limited. In the unlimited sampling mode, subjects could make as many switches and check each stimulus for as many times as they needed.

The motor effort during the action stage took the form of moving the cursor to the corresponding choice button and clicking on it to report the final choice. The choice buttons were two small white squares displayed on the lower half of the screen, vertically 7 cm from the centers of the stimuli. We set two types of trials differentiated by the location of the choice buttons: In the first type, the buttons were horizontally centered, so the motor effort (measured by the moving distance) to drag the cursor from the two stimuli to the buttons was approximately the same. In the second type, the buttons were placed under the right stimulus, so that the required motor effort would be less if the subject sampled the right stimulus last and started from there to reach for the buttons.

The display screen size was 28.5 × 18 cm, resolution 1280 × 800 pixels, refresh rate 60 Hz. The screen was placed 50 – 70 cm in front of the subjects. System mouse acceleration was disabled to make the cursor movement on the screen linearly map the actual movement of the mouse. Subjects were told not to pick up the mouse from the surface of the desk amid each trial. Mouse trajectory was recorded from the moment the trial was triggered to when a button was clicked (sampling rate 60 Hz). We also recorded the final decision in each trial. The stimuli and mouse tracking codes were programmed in MATLAB Psychtoolbox-3.

### Participants and procedure

A total of 24 subjects participated in the study (13 females, age 20 – 30); all of them were university students. Subjects wore glasses for vision correction if needed. The research was approved by the institutional ethics committee of Eotvos Lorand University, Hungary. All subjects provided informed written consent, and none declared any history of neurological diseases.

To avoid previous experimental processes interfering with later sampling patterns, we divided the subjects into 3 groups, each containing 8 subjects, and each group of subjects only performed in a single experimental condition (Table 1). After 10 practice trials to get familiar with the paradigm, each subject performed 2 blocks of 60 trials. A short break (5 – 10 minutes) took place between the blocks. The complete experiment took approximately 40 – 60 minutes per subject.

**Table 1.**
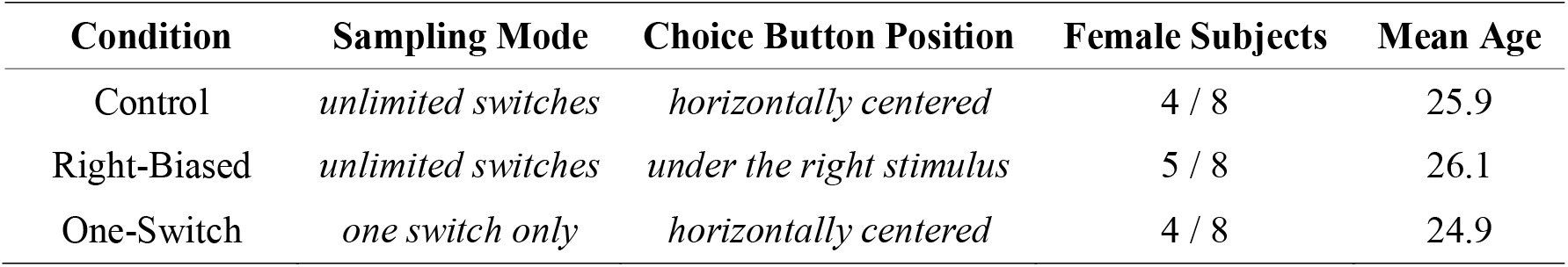
Details of the experimental condition settings.

### Data analysis

#### Sampling patterns

*Decision time* for a trial was defined as the elapsed time from the onset of the stimuli to when the final decision was made, excluding the time of action execution. The dividing line between the sampling stage and the action stage was the moment when a downward y-axis component of the cursor velocity exceeded the threshold.

*Horizontal moving distance* during sampling was defined as the total moving distance of the cursor on the screen along the x-axis within the sampling stage of a trial.

To test the linear relationship between the variables depicting sampling patterns, we performed linear mixed-effects regressions with random effects for subject-specific intercepts and slopes.

#### Psychometric curves

Psychometric curves were fitted to the data pooled across all subjects within each group or all simulation trials in the same condition using the generalized linear model (GLM) with the logit link function.

#### Comparing lines and curves

To compare two regression lines, we used a generalized ANCOVA allowing different slopes and intercepts:

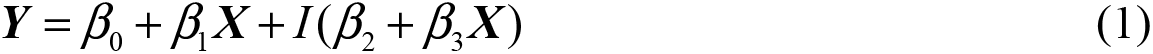

where *β*_0_, *β*_1_, *β*_2_ and *β*_3_ were free parameters, ***X*** was the predictor variable, and *I* was the indicator variable whose value was 0 for the reference group and 1 for the other group.

To compare two psychometric curves, we fitted the data to the following logistic function:

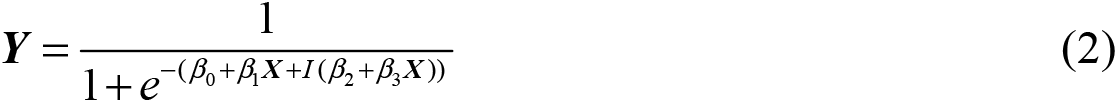

Then, we tested the null hypotheses *β*_2_ = 0 and *β*_3_ = 0 with the two-tailed one-sample t-test to compare the intercepts and slopes (steepness) of the two curves.

### Bayesian Network modeling

We listed all three possible Bayesian Network models for different hypotheses regarding the causal relationship between the decision variable before the last sampling, the last sampled item and the final choice in the trial. The conditional probability of choosing the right item given that it is last sampled was calculated under each hypothesis and compared with the empirical results.

For mathematical details of the models, see S1 Supporting Information.

To calculate the conditional probability *p*(right chosen | right last sampled) from the behavioral data, we first fitted a psychometric choice curve (probability of choosing the right item vs. difference between the proportions of white dots in the stimuli) to the trials in which the right item was sampled last for each subject individually, and then marginalized the difference between the stimuli. The mean *p*(right chosen | right last sampled) across the subjects in each group was compared with the value 0.5 (the probability without bias) using the one-tailed one-sample t-test. The One-Switch group and the Right-Biased group were compared with the Control group using Dunnett’s test after a one-way ANOVA.

### aDDM simulation

We built our aDDM following Krajbich et al. [15]. We set the relative value (*r*_left_ and *r*_right_) to the proportion of white dots in each stimulus. The range of *r*_left_ and *r*_right_ in the experiment was [0.25, 0.75]. The decision variable (*DV*) started from 0 in each simulation trial, and the decision barriers were −1 for the left stimulus and +1 for the right stimulus. We applied the multiplicative model [22]. The drift rates (*v*) in the model were defined as:

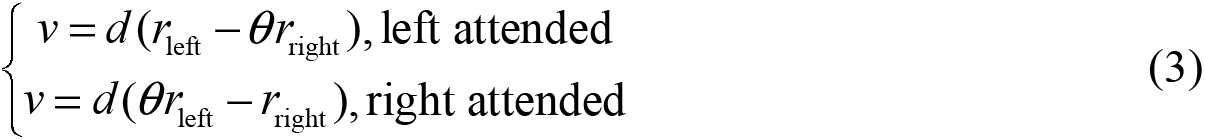

where *d* was the value scaling parameter, and *θ* was the multiplicative attentional discounting parameter. Specifically, during the first sampling, the unattended stimulus was assigned a mean value *r*_mean_ = 0.5 instead of the real value because the subject had not sampled that stimulus yet. Let *DV*_*t*_ denote the value of the decision variable at time *t*. For every time step Δ*t*,

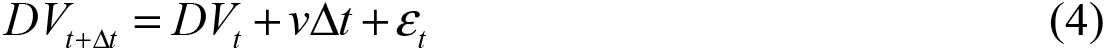

where *ε*_*t*_ was drawn from a zero-mean Gaussian distribution with standard deviation *σ*. We assumed that the first sampling falls on the left stimulus with a fixed probability (Control: 0.59, One-Switch: 0.57, from empirical data), its duration drawn from a fixed gamma distribution. Each successive sampling epoch fell alternatively on the left and right stimulus and would continue until it reached a max time limit drawn from another fixed gamma distribution or until the decision variable reached one barrier. The parameters of the two gamma distributions were fitted with maximum likelihood estimation (MLE) to the empirical sampling time data in the Control condition. Time step Δ*t* was set to 10 ms. For human subjects in the Control group, the max number of switches in a single trial was 10, so we discarded simulations with more than 10 switches.

We fitted the three parameters in the model (*θ*, *d* and *σ*) to the empirical data pooled across all subjects: For each set of parameters, we ran a fixed number of valid simulations (240 for the coarse search and 960 for the finer search) and compared the results with behavioral data using the following error metric:

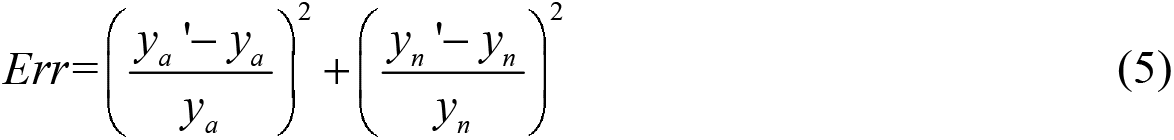

where *y*_*a*_ = 0.9042 and *y*_*n*_ = 2.0698 were the accuracy and the mean number of switches calculated from the 960 trials pooled across the 8 subjects in the Control group, while *y*_*a*_’ and *y*_*n*_’ were the accuracy and the mean number of switches across all simulations. We performed a grid search for the best fitting parameters: In the *i*-th iteration, we tested the parameter sets given by the cross product of {*θ*_*i*1_, *θ*_*i*2_, *θ*_*i*3_}, {*d*_*i*1_, *d*_*i*2_, *d*_*i*3_} and {*σ*_*i*1_, *σ*_*i*2_, *σ*_*i*3_}. Let (*θ*_*i*_, *d*_*i*_, *σ*_*i*_) denote the parameters that generated the smallest *Err* value, then in the (*i*+1)-th iteration we tested a finer grid given by the cross product of {*θ*_*i*_ – 0.5Δ*θ*_*i*_, *θ*_*i*_, *θ*_*i*_ + 0.5Δ*θ*_*i*_}, {*d*_*i*_ – 0.5Δ*d*_*i*_, *d*_*i*_, *d*_*i*_ + 0.5Δ*d*_*i*_} and {*σ*_*i*_ – 0.5Δ*σ*_*i*_, *σ*_*i*_, *σ*_*i*_ + 0.5Δ*σ*_*i*_}, where Δ*θ*_*i*_, Δ*d*_*i*_ and Δ*σ*_*i*_ were the step sizes used in the *i*-th iteration. The initial values were {0.1, 0.5, 0.9} for *θ*, {0.001, 0.005, 0.009} for *d*, and {0.01, 0.05, 0.09} for *σ* in the coarse search and {0.6, 0.7, 0.8} for *θ*, {0.004, 0.005, 0.006} for *d*, and {0.03, 0.04, 0.05} for *σ* in the finer search. We stopped the iterations when the step sizes became smaller than 0.5% of the parameter values. The final fitting results were *θ* = 0.67, *d* = 0.0051 and *σ* = 0.038.

For the One-Switch condition, we used the same set of parameters (*θ*, *d* and *σ*) in the Control condition, but only two sampling epochs were allowed. The second sampling would continue until the decision variable reached one barrier. We discarded the simulations in which the second sampling exceeded 3000 ms, which was the max duration of the second sampling for 99.5% empirical trials in the One-Switch condition. The decision time for the simulations was calculated by adding the mean transition time (delay between the sampling epochs) measured from behavioral data to the total time length of the sampling epochs in the simulations.

We simulated the model for 960 valid trials (the same sample size as the empirical data pooled across all subjects in each condition) using the best fitting parameters above for the Control condition and the One-Switch condition separately and compared the results with human behavioral data.

## Results

### General sampling patterns

Firstly, we studied the general sampling patterns in the Control condition. We plotted the horizontal mouse cursor position recorded during the sampling stage against elapsed time in each trial. Fig 2 shows the time series of the cursor position from a single block performed by one subject: The 60 trials in the block were sorted by the start position. The horizontal positions between the two stimuli were linearly mapped to [0, 1] and shown in a red-blue color scale. The typical sampling pattern was to switch the cursor once or multiple times between the two stimuli. The cursor paused mostly at either the leftmost or the rightmost, meaning that only one of the stimulus was clearly visible at a time. Therefore, we can assume that the eye gaze and the attention of the subject switched between the stimuli together with the cursor, which enables the comparison between our paradigm and former sequential sampling tasks and models.

**Fig 2.**
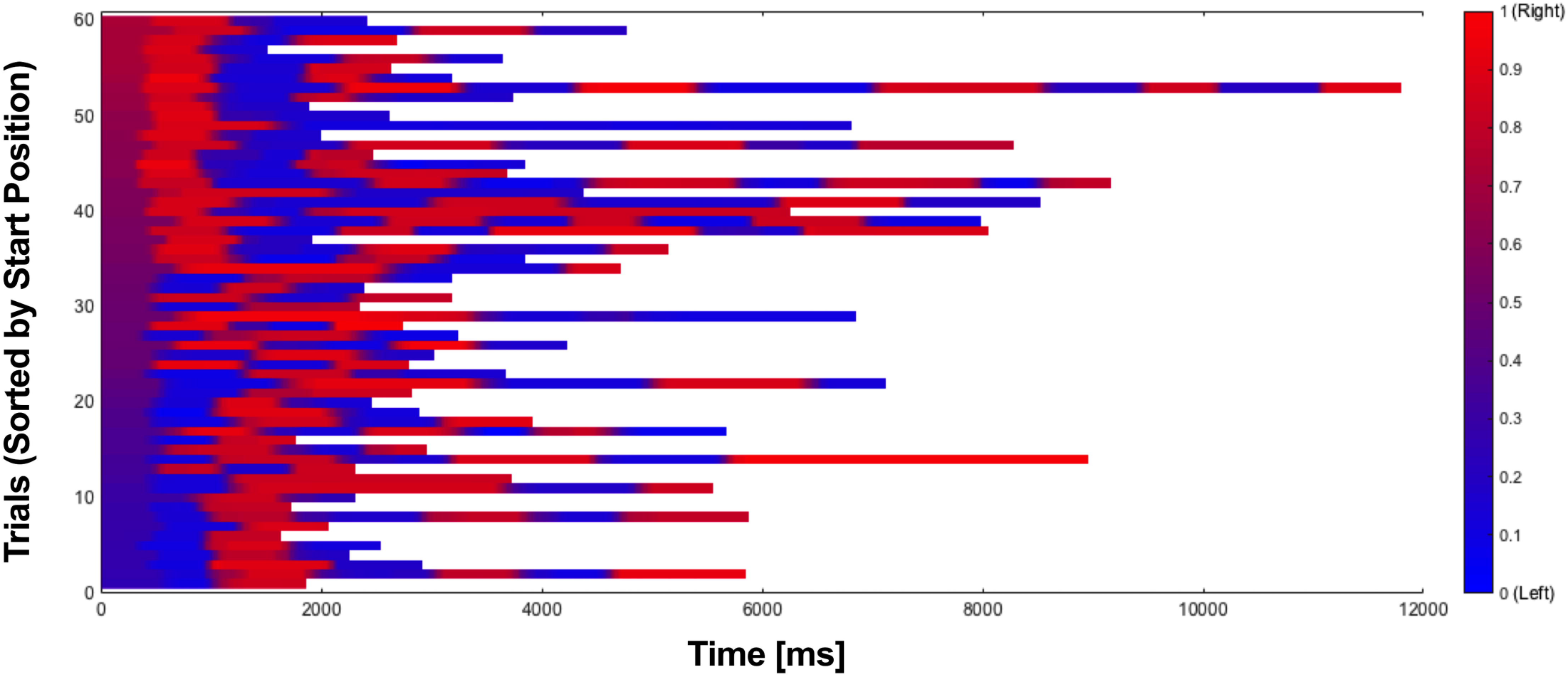
Typical time series of the horizontal mouse cursor position during sampling. Data were from a single block (60 trials) performed by one subject in the Control group and sorted by the start position in each trial. Red color indicates that the current cursor position is closer to the right stimulus, while blue indicates that the cursor is closer to the left.

If a subject made *n* switches in a trial, there would be *n*+1 sampling epochs alternatively assigned to the two stimuli. Assuming that each sampling period has approximately the same duration, the decision time should linearly correlate with the number of switches in each trial. Moreover, most of the motor effort during sampling was spent on switching the cursor from one stimulus to the other, the distance between them fixed. Therefore, the total motor effort within a trial (measured by the horizontal moving distance of the cursor) should also linearly correlate with the number of switches. Fig 3A shows the histogram of the number of switches made in all 960 trials performed by subjects in the Control group: In 42.5% trials only one switch was made, and the percentage of the trials decreased as the switches made in them increased. Fig 3B and 3C show that the decision time (linear mixed-effects regression: *slope* = 1276.6 ms, *P* = 2.8×10^−21^; Pearson’s *r* = 0.78) and the horizontal moving distance (linear mixed-effects regression: *slope* = 17.8 cm, *P* = 2.6×10^−54^; Pearson’s *r* = 0.90) linearly correlated with the number of switches.

**Fig 3.**
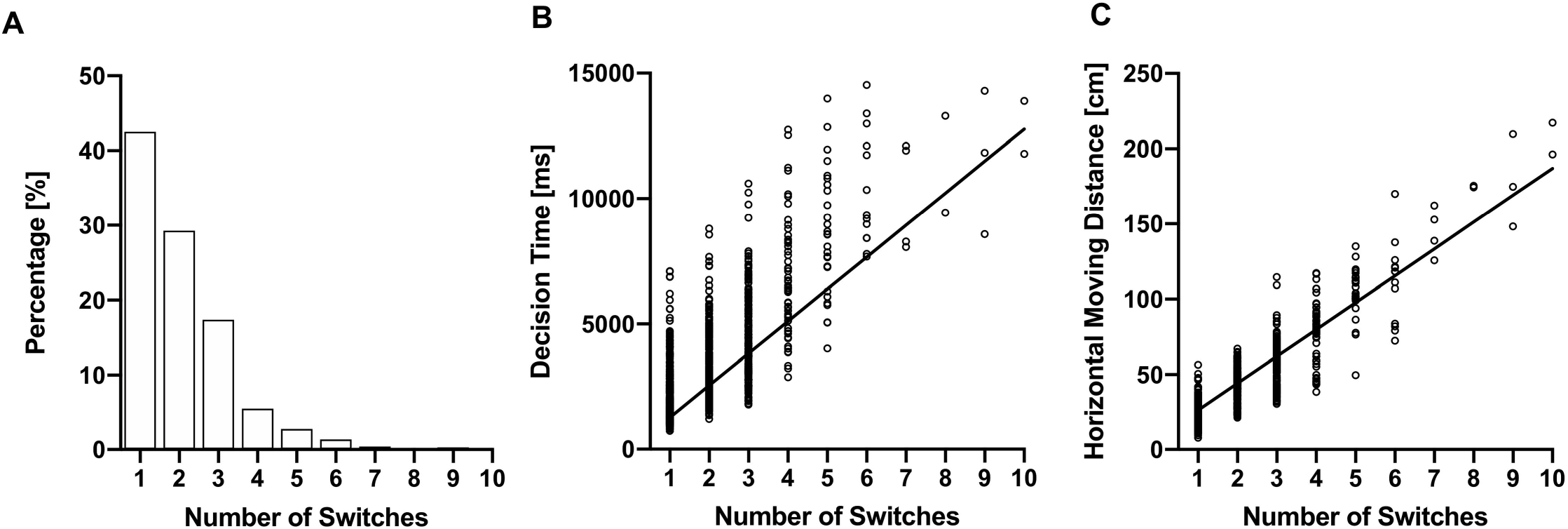
Number of switches within each trial, and its correlation with decision time and horizontal moving distance. **(A)** Histogram of the number of switches. **(B)** Linear correlation between decision time and number of switches during the trials. Mixed-effects regression: *slope* = 1276.6 ms, *P* = 2.8×10^−21^; Pearson’s *r* = 0.78. **(C)** Linear correlation between horizontal moving distance and number of switches during the trials. Mixed-effects regression: *slope* = 17.8 cm, *P* = 2.6×10^−54^; Pearson’s *r* = 0.90. Data included 960 trials pooled across 8 subjects in the Control group.

### Influences of motor effort on sampling patterns

The total motor effort within one single trial consisted of three parts: first, to drag the cursor from the start position to the first sampled item; second, to switch between the items one or more times during sampling (each switch took approximately the same moving distance, as discussed previously); third and lastly, to drag the cursor from the last sampled item to the choice buttons. We studied how motor cost in the different parts interacted with the decision-making process (Fig 4):

**Fig 4.**
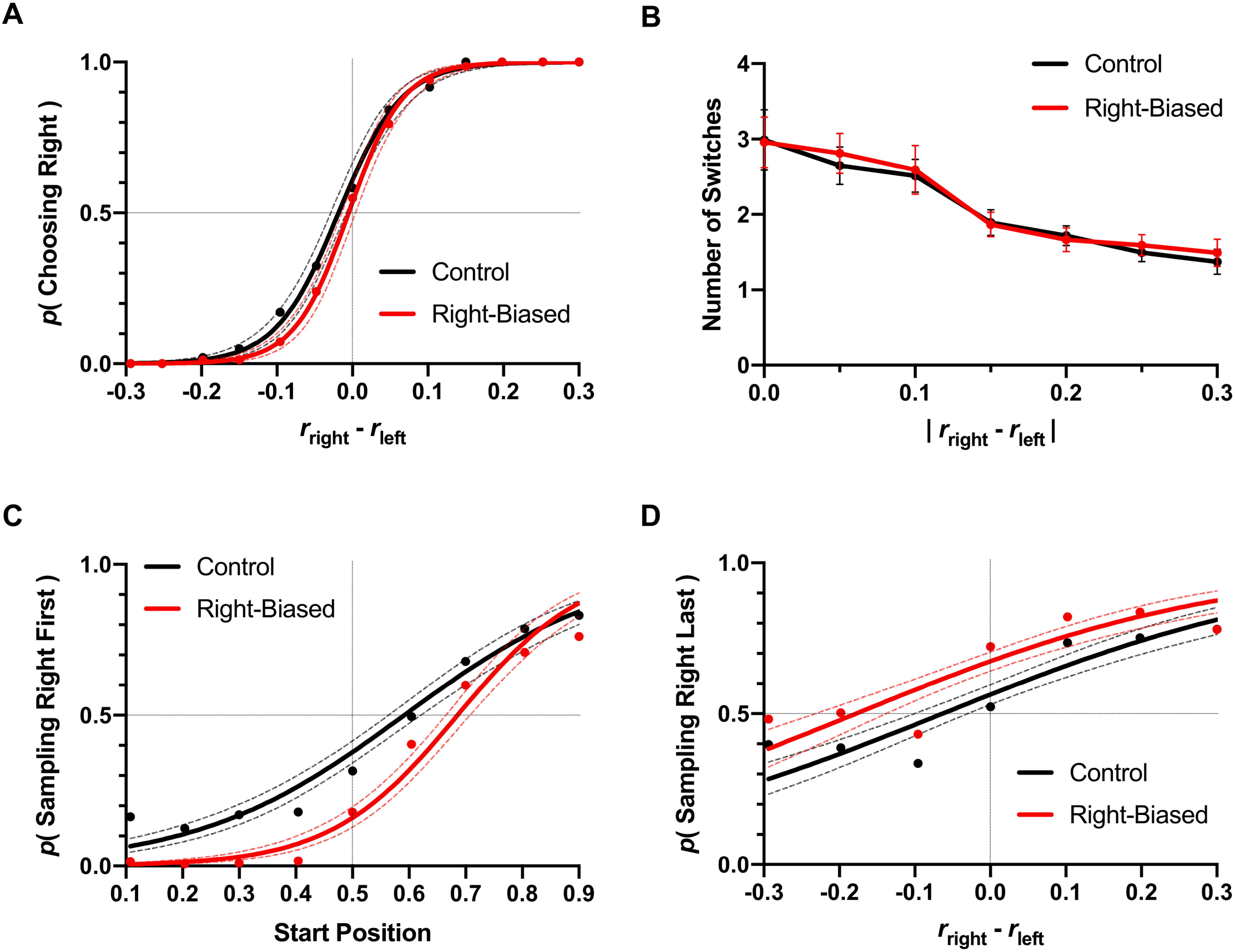
Right-Biased condition vs. Control condition. **(A)** Psychometric choice curves. **(B)** Number of switches against trial difficulty, as measured by the absolute difference between the proportions of white dots in the stimuli. **(C)** Psychometric curves for the first sampled item. **(D)** Psychometric curves for the last sampled item. Error bars show 95% confidence intervals. Data included 960 trials pooled across 8 subjects in each condition.

Fig 4A shows the psychometric choice curves in the Control and the Right-Biased conditions: There was no statistically significant difference between the two curves (intercept: *P* = 0.0958; slope: *P* = 0.1358), and the overall accuracy was also similar (Control: 90.4%; Right-Biased: 91.5%; unpaired two-tail t-test between individual subjects in the two groups: *P* = 0.4599). The difference in motor costs during the action phase did not bias the decisions of the subjects in our experimental paradigm. One possible reason is that the difference was not directly related to the final choice; another possibility is that explicit knowledge of the motor cost would help to avoid integrating irrelevant factors into the decision to maintain high accuracy [23].

We plotted the number of switches made in the trials against trial difficulty (measured by the absolute difference between the proportions of white dots in the stimuli) in Fig 4B: In both conditions, the number of switches decreased with trial difficulty (significant slopes in mixed-effects regression: *P* = 4.8×10^−10^ for Control and *P* = 1.9×10^−8^ for Right-Biased). There was no significant difference between the two conditions (intercept: *P* = 0.5216; slope: *P* = 0.8516). The motor cost during sampling correlated with the number of switches, therefore we concluded that the more difficult the trials were, the more motor effort would be invested into the sampling process.

Next, we examined the influences of motor cost upon the first and the last sampled items. Fig 4C shows that the start position and the choice button position both affected the first sampled item: In the Control condition, subjects tended to sample the item closer to the start position, which would reduce the first part of the total motor cost. However, we observed a systematic bias to sample the left item first, which may be related to the cultural habit of dealing with items in left-to-right order (for example, people usually read from left to right). In the Right-Biased condition, subjects showed an extra tendency to go for the left item first (significantly different intercepts: *P* = 2.9×10^−9^). The subject was most likely to make only one switch (Fig 3A); in that case, starting from the left item would lead to taking the shorter path from the right item to the buttons, reducing the last part of the motor effort. In Fig 4D we plotted the probability of sampling the right item last against the difference between proportions of white dots in the two stimuli: Generally, subjects were more likely to sample the stimulus with more white dots last. In the Right-Biased condition, subjects preferred to sample the right stimulus last (significantly different intercepts: *P* = 2.9×10^−6^), which would reduce the motor cost during the action phase.

### Influences of the decision variable on sampling patterns

The last-sampling bias is the phenomenon that subjects are more likely to choose the last sampled stimulus. Such a bias has been reported in several human decision-making studies of both preferential and perceptual decisions [15, 17]. However, the causal relationship behind the last-sampling bias is not completely clear: Do subjects tend to choose a stimulus because it is the last sampled one, or do they tend to sample the particular stimulus last because they already want to choose it, or both? According to the aDDM, the evidence accumulation rate for the stimulus not being sampled is discounted, so the decision variable will be more likely to reach the barrier at the last sampled side [15]. Otherwise, the aDDM assumes that the current decision variable has no backward influence upon sampling patterns. There is rare evidence supporting that the temporary decision has a causal effect on the allocation of attention during sampling [19].

### Bayesian Network modeling

To study the causal relationship between the last sampled item and the decision, we built a Bayesian Network model quantifying the size of the last-sampling bias. Fig 5 displays the graphical models for the networks: Naturally, the final decision depends on the decision variable. In the One-Switch condition, the last sampled item is the alternative of the first sampled one, which in turn depends on the motor cost measured by the distance from the start position to the stimulus. In the Right-Biased condition, the last sampled item depends on the motor cost in the action stage. Apart from the common dependency structures described above, there are three possible models with different hypotheses on whether the last sampled item depends on the decision variable and whether the final choice depends on the last sampled item:

**Fig 5.**
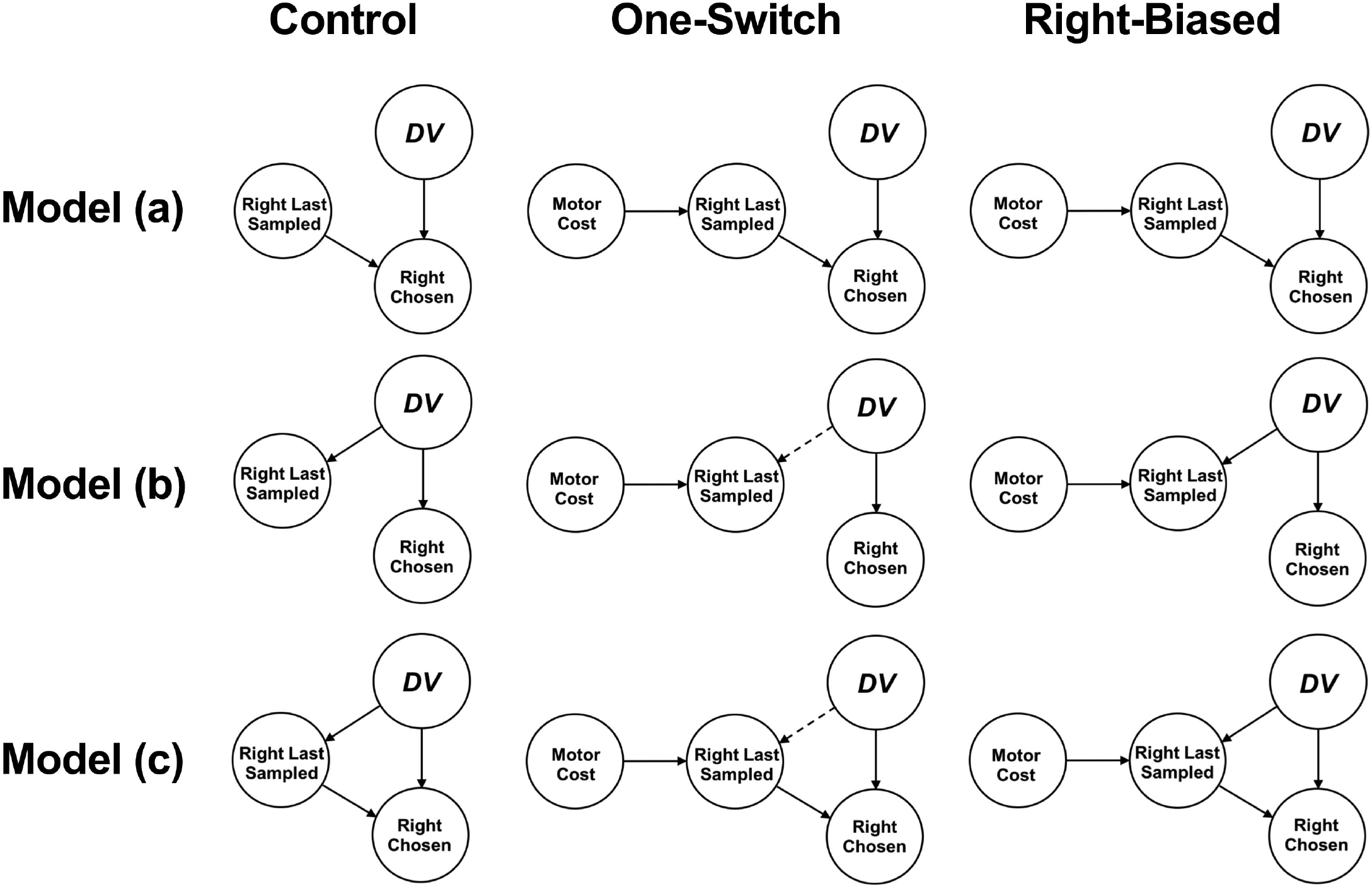
Causal relationship for the last-sampling bias. Graphical models for possible hypotheses regarding the conditional dependency relationship between the decision variable before the last sampling, the last sampled stimulus and the final choice. Arrows show dependency between the events or variables, and dashed arrows show dependency assumed to exist generally but absent in the specific experimental condition.

#### Model (a)

Like the aDDM, the first model assumes that the final decision depends on the last sampled item, but the last sampled item is independent of the decision variable. Therefore, in the Control condition we have:

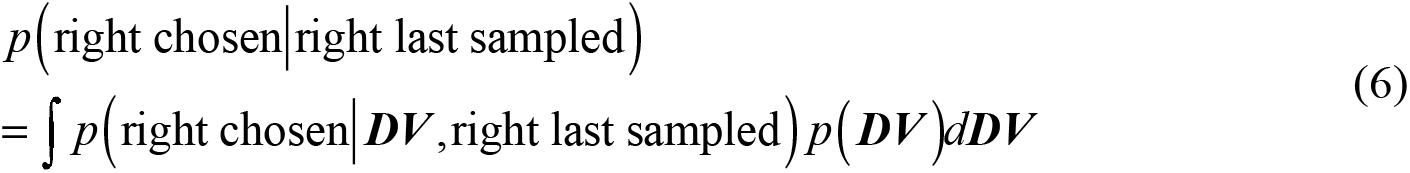

where ***DV*** is the value of the decision variable exactly before the last sampling epoch starts. *p*(right chosen | right last sampled) measures the size of the last-sampling bias. The bias is due to the term *p*(right chosen | ***DV***, right last sampled), which can be regarded as a function of ***DV***: For each given value of ***DV***, *p*(right chosen | ***DV***, right last sampled) > *p*(right chosen | ***DV***). In the aDDM, *p*(right chosen | ***DV***, right last sampled) is the probability for the decision variable to drift to the right boundary at the end of the last sampling, decided by the aDDM parameters (*d*, *σ* and *θ*) and the relative values for the two stimuli.

In the One-Switch condition, the dependency structure between ***DV***, the last sampled item and the final choice remains the same, but the following analysis explains why the size of the last-sampling bias will increase (see our aDDM simulation results): At the beginning of each trial, the decision variable is set to 0; as the sampling time elapses, more drift steps (*v*Δ*t* + *ε*_*t*_) are added to the decision variable, so its variance increases. When only two sampling epochs are allowed, the elapsed time before the last sampling is shorter, thus *p*(***DV***) will have a narrower variance. In that case, the product of a biased *p*(right chosen | ***DV***, right last sampled) and *p*(***DV***) will be larger than that in the Control condition.

In the Right-Biased condition, the value of *p*(right chosen | right last sampled) will not change because no term in Equation (6) depends on the motor cost to reach for the buttons.

#### Model (b)

On the contrary, the second model assumes that the last sampled item depends on ***DV***, but the final decision is independent of the last sampled item. Therefore, in the Control condition,

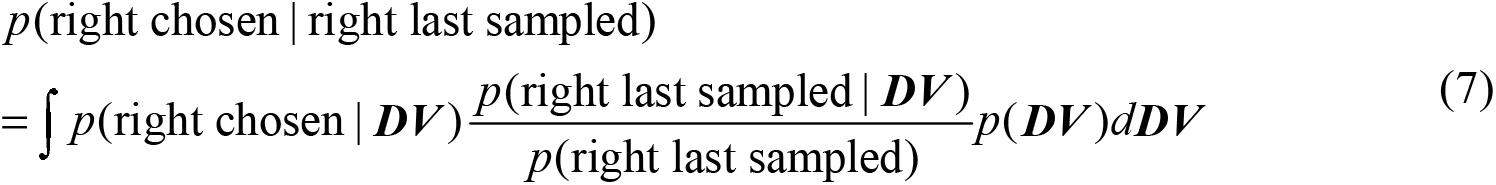

Under that hypothesis, the last-sampling bias is due to the term *p*(right last sampled | ***DV***), which is also a function of ***DV***: When ***DV*** > 0, *p*(right last sampled | ***DV***) > *p*(right last sampled); when ***DV*** < 0, *p*(right last sampled | ***DV***) < *p*(right last sampled).

In the One-Switch condition, the last sampled item no longer depends on ***DV***, so:

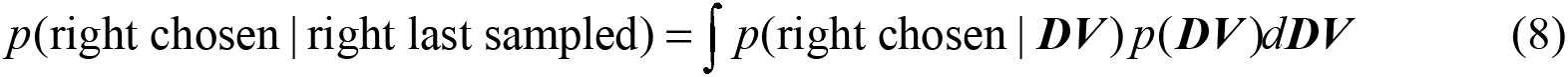

Thus *p*(right chosen | right last sampled) will fall back to 0.5.

In the Right-Biased condition, subjects tend to sample the right item last. We assumed that such a tendency is independent of ***DV***, so the probability *p*(right last sampled | ***DV***) and *p*(right last sampled) will rise by the same additive amount. Compared with the Control condition, the term *p*(right last sampled | ***DV***)*p*(right last sampled)^−1^ will become smaller when ***DV*** > 0 and larger when ***DV*** < 0, resulting in a decreased size of the last-sampling bias.

#### Model (c)

This model shows a third possibility that the last sampled item depends on ***DV***, and the final decision also depends on the last sampled item. In the Control condition:

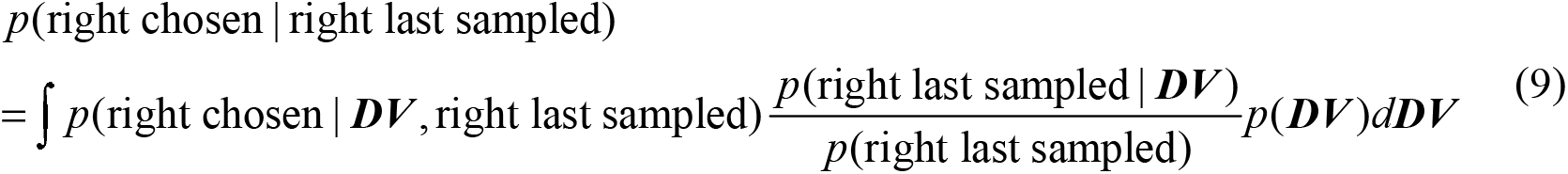

In Equation (9), the total bias in *p*(right chosen | right last sampled) has two sources: *p*(right chosen | ***DV***, right last sampled) and *p*(right last sampled | ***DV***). The last sampled item is more likely to be chosen, and the temporarily winning item is more likely to be sampled last.

Under such assumptions, the last-sampling bias should remain in the One-Switch condition because the term *p*(right chosen | ***DV***, right last sampled) is biased, but the size will decrease because the term *p*(right last sampled | ***DV***) now disappears.

In the Right-Biased condition, *p*(right chosen | right last sampled) will also become smaller similar to that in Model (b).

### Model predictions vs. empirical results

Let *p*_Control_, *p*_One-Switch_ and *p*_Right-Biased_ denote *p*(right chosen | right last sampled) in each specific experimental condition. We summarized different model predictions and the empirical results in Table 2: Among the three hypotheses, only Model (c) correctly predicted the behavioral data. Therefore, we concluded that the causal relationship between sampling patterns and the decision is bidirectional.

**Table 2.**
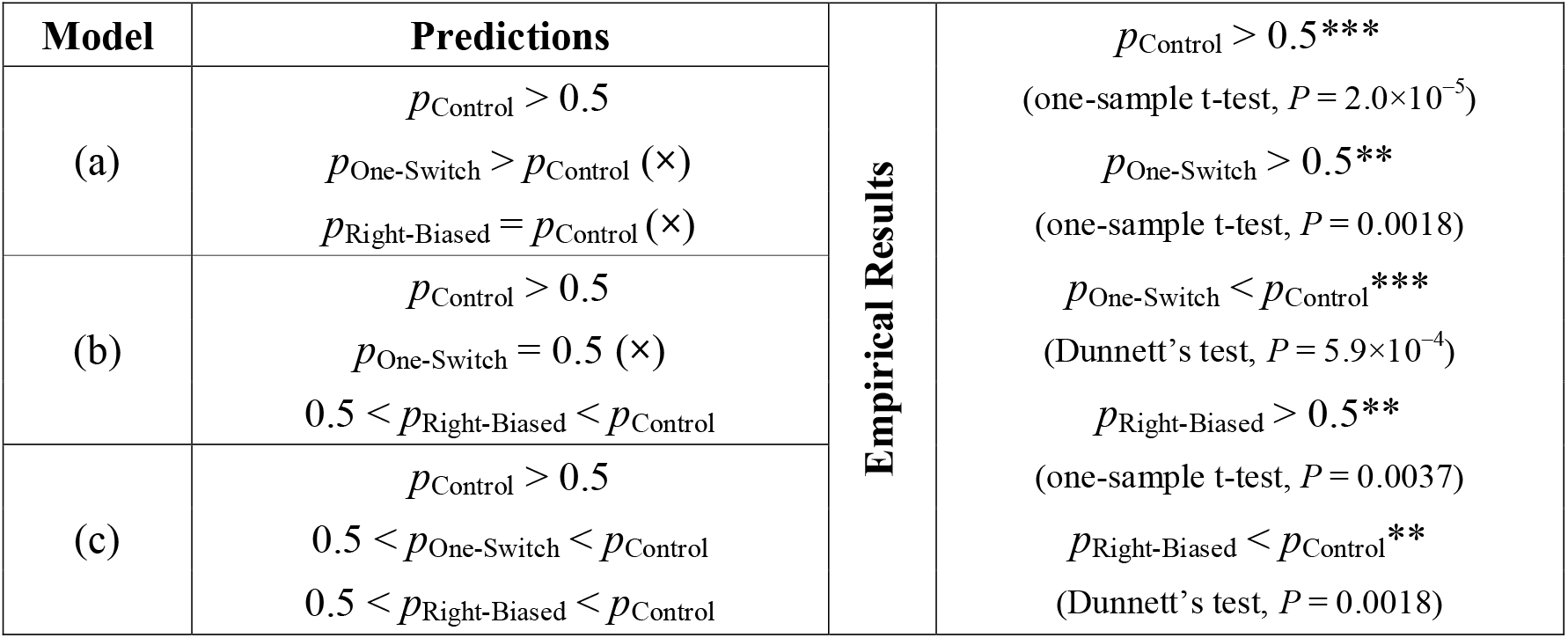
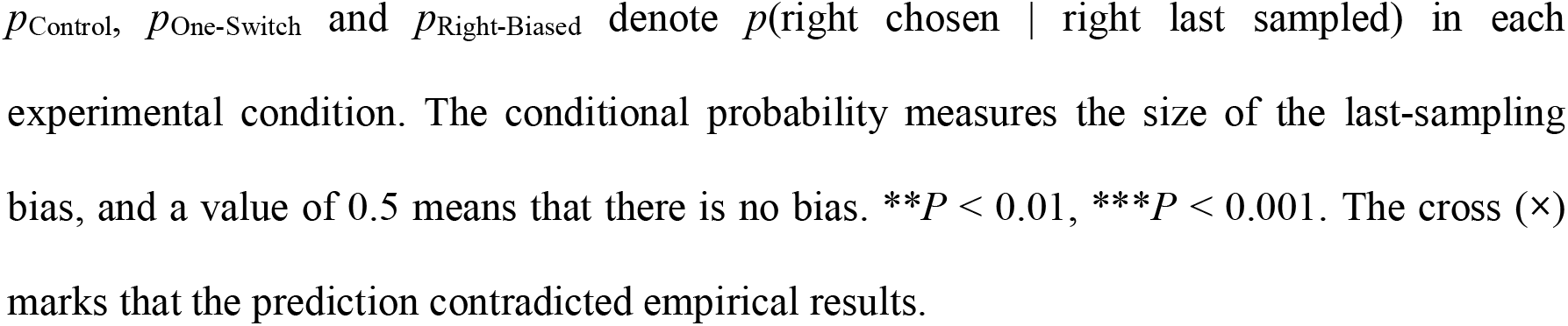
Summary of different model predictions about the last-sampling bias and the empirical results.

### aDDM simulation

On top of the theoretical Bayesian Network analysis, we also ran an aDDM simulation to test whether the decision variable can feedback into the sampling patterns. In our simulations, each sampling epoch was focused alternatively on the two stimuli until it reached a time limit randomly drawn from a distribution fitted to the empirical data. In the One-Switch condition, only one switch of attention was allowed. Each simulation ended when one of the decision boundaries was reached. Therefore, the allocation of attention in the aDDM was independent of the current decision variable. Fig 6 shows the comparison between simulated and empirical results:

**Fig 6.**
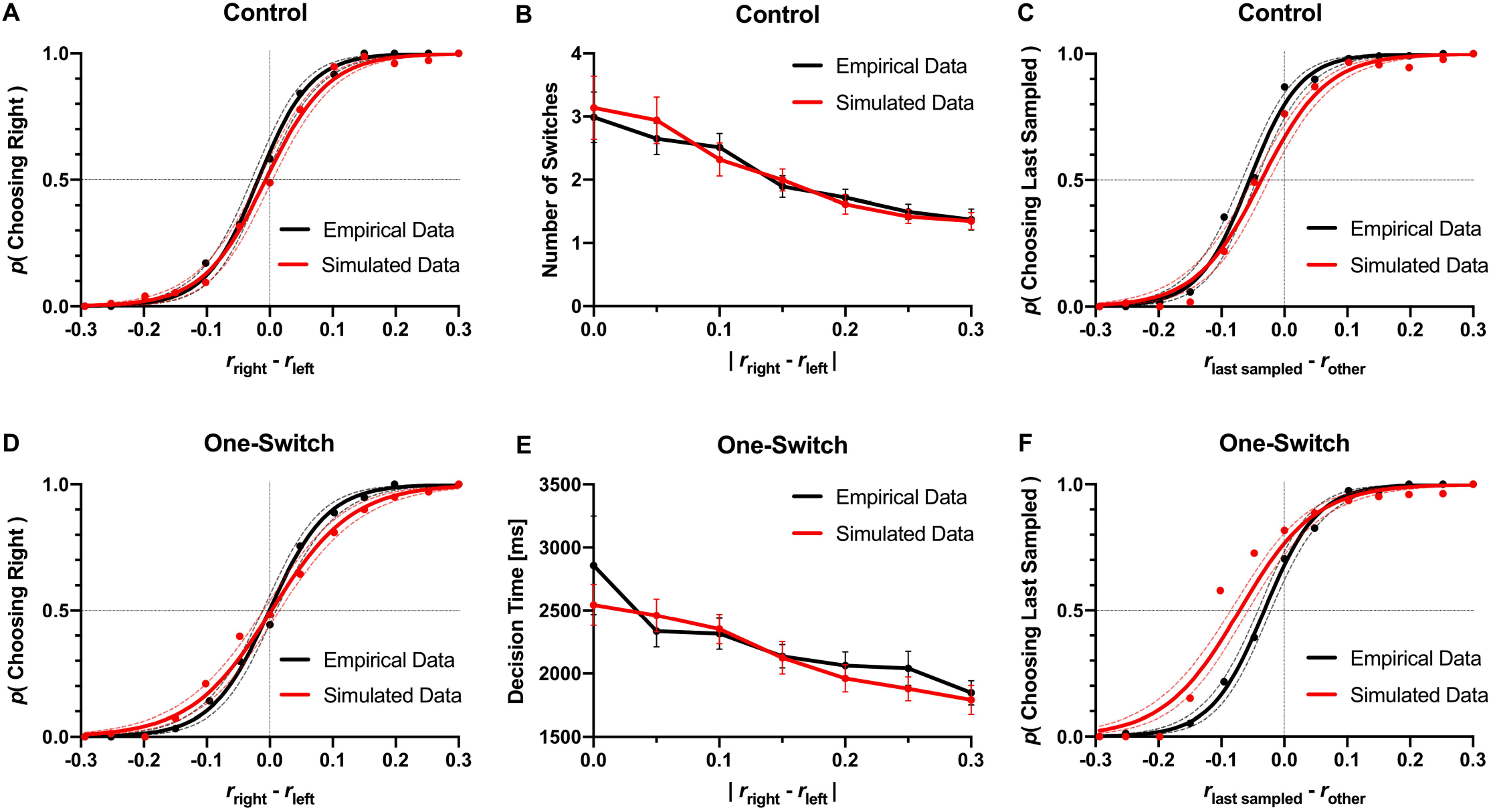
aDDM simulation results vs. empirical data. **(A)** Psychometric choice curve in the Control condition. **(B)** Number of switches in the Control group against trial difficulty, as measured by the absolute difference between the proportions of white dots in the stimuli. **(C)** Last-sampling bias in the Control group. The horizontal axis shows the difference between the proportions of white dots between the last sampled stimulus and the other, while the vertical axis shows the probability of choosing the last sampled stimulus. **(D)** Psychometric choice curve in the One-Switch condition. **(E)** Decision time in the One-Switch group against trial difficulty. **(F)** Last-sampling bias in the One-Switch group. Error bars show 95% confidence intervals. Empirical data included 960 trials pooled across 8 subjects in each condition; simulated data included 960 trials in each condition.

Firstly, we compared the psychometric choice curves: There was no statistically significant difference between the psychometric choice curves for the simulations and the human subjects (intercept: *P* = 0.0679; slope: *P* = 0.0610) in the Control condition (Fig 6A). In the One-Switch condition, there was no significant difference between the intercepts of the curves (*P* = 0.7116), but the slope (steepness) for the simulated data was significantly smaller than that of the empirical data (*P* = 3.2×10^−4^), meaning that the overall accuracy in the simulations was lower (Fig 6D). When only one switch was allowed, the choice accuracy of the simulations reduced from 90.1% to 84.4%, while for the human subjects there was no significant reduction (unpaired one-tail t-test between individual subjects in the two groups: *P* = 0.8082).

Next, we compared the number of switches in the Control condition (Fig 6B): There was no statistically significant difference between the simulated and empirical results regarding the number of switches against trial difficulty (intercept: *P* = 0.2690; slope: *P* = 0.2384). In the One-Switch condition, we compared the decision time instead of the number of switches (for it is constantly 1): There was no significant difference between the simulated and empirical results regarding the decision time against trial difficulty ((Fig 6E, intercept: *P* = 0.6125; slope: *P* = 0.2257).

Finally, we focused on the last-sampling bias: In Fig 6C and 6E, we plotted the probability of choosing the last sampled item against the difference between the proportions of white dots between the last sampled stimulus and the other. All the curves had an intercept larger than 0.5, showing a tendency to choose the last sampled item, but the sizes of the bias were different: The curve intercept for the simulations was significantly lower than empirical (*P* = 5.8×10^−4^) in the Control condition and significantly higher than empirical (*P* = 0.0166) in the One-Switch condition. Denying that subjects would switch back to sample the winning item but assuming random switches all the while, the aDDM underestimated the last-sampling bias in the Control condition and overestimated it in the One-Switch condition. The simulation results matched our Bayesian Network analysis and implied that the current decision had a causal effect on the sampling patterns.

## Discussion

In summary, the adaptive sampling behavior during perceptual decision-making exhibited the following patterns: First, the number of switches between the alternatives correlated with the difficulty of the task: the more difficult the task was, the more times the stimuli were resampled. Second, the sampling sequence was decided considering the start position and the choice button position in an attempt to minimize the total motor effort. Third, attention was biased to the eventually chosen item during the last sampling epoch. Combining the modeling results, we concluded that both motor cost and the temporary decision have a causal influence upon the pattern of attention allocation during sampling.

Having reviewed recent computational models, behavioral studies and neural recording results, Wispinski et al. [24] concluded that decision-making is a continuous process from the presentation of behaviorally relevant options until movement completion. Previous studies suggested that motor effort related to the action phase can influence the decision [4–6, 8], and our results provided an extended conclusion that sampling behavior was also influenced by the motor effort in different stages of the decision-making process. It supported the idea that sensorimotor aspects should be considered as an actively integrated part of the decision-making process. Further, several studies focused on the representation of motor effort and how it is related to cost minimization in decision-making as well as motor control [25, 26]; future studies may quantify the effect of motor cost on sampling behavior with similar methods.

The relationship between attention and eye movement during decision-making has been studied abundantly [27], but researches highlighting limb and body movements during the sampling process are rare, even though in naturalistic circumstances such movements usually cooperate with eye movements to sample relevant information better. In our research, we designed a paradigm based on computer mouse tracking in which both gaze shift and hand movement (moving the mouse) were necessary to switch attention between the options. Although mouse tracking and eye tracking are both commonly applied process tracking methods in decision-making research, their original purposes are slightly different: While eye tracking mostly target on attention and information searching strategies, mouse cursor tracking data reflect more about indecision and momentary preference [28]. In our paradigm, however, subjects must move the cursor closer to get a better view of each stimulus, as if approaching a real object to have a better look. In this way, the mouse trajectory can reflect attention during sampling as eye traces did in previous studies. Moreover, our paradigm can be applied to study eye-hand cooperation and coordination during decision-making as well.

Traditionally, sequential sampling models assume that during decision-making, subjects sample their options continuously until the relative evidence for one option reaches a predetermined threshold, and such models capture the speed-accuracy trade-off phenomenon well [19, 29, 30]. Interestingly, our results showed that subjects would make extra sampling epochs during which the accuracy of the decision has not been improved significantly. One possible explanation is that subjects were switching back to the previously sampled stimuli again to verify their preliminary decision [31]. Similar to other studies [18, 32, 33], we observed an attentional bias to the finally chosen option during the later sampling epochs. Mullett and Stewart [32] suggested that such a bias may be due to a relative instead of absolute stopping rule. According to Krajbich [19], even as the decision variable evolves and one option emerges as the winning one, it is still optimal to continue sampling information randomly instead of favoring the leading option, since the information from both the winning and losing options are of equal importance. However, the accuracy of the decision will not necessarily decrease if the attentional bias happens at the later stage of sampling when the main task is to validate the decision. This validating phase may be longer for perceptual decisions, for people tend to respond with more caution in perceptual decisions than in preferential decisions, especially when the stimuli are ambiguous [34]. Meanwhile, how the sub-thresholds within the preliminary decision phase and the validating phase are determined remains to be discussed.

Finally, our study provided evidence for the bidirectional causal relationship between attention and decisions by Bayesian Network modeling. Bayesian Networks have been customarily applied for probabilistic causal dependence assessment and inference in a wide range of areas [35], including life science researches [36, 37]. It is capable of depicting and predicting the conditional dependences between experimental variables through observed data, thus becomes a beneficial tool for psychological studies. In our study, we listed all possible network structures corresponding to different hypotheses on the causal relationship between the last sampled item, the decision variable and the chosen item. Then, we compared the predicted conditional probability of choosing the last sampled item with empirical data. Contrary to previous literature [19], our results imply that rather than randomly switching between the options, attention is drawn to the winning item during sampling. This finding may lead to some modification to the basic assumptions of the aDDM in the future.

## Supporting information

S1 Supporting Information. Mathematical details of the Bayesian Network modeling.

S2 Video. Demo trials recorded from the screen.

## Acknowledgements

We would like to express our sincere gratitude to Prof. László Mérő’s supervision as well as his support and encouragement. We are also grateful for the help on practical issues from the Faculty of Education and Psychology ELTE and the kind participation of all the subjects in the experiments.

## Supporting information

**S1 Supporting Information. Mathematical details of the Bayesian Network modeling.**

**S2 Video. Demo trials recorded from the screen.**

## References

1. Lepora NF, Pezzulo G. Embodied choice: how action influences perceptual decision making. PLoS computational biology. 2015 Apr 7;11(4):e1004110. doi: 10.1371/journal.pcbi.1004110

2. Connors BL, Rende R. Embodied Decision-Making Style: Below and Beyond Cognition. Frontiers in psychology. 2018;9. doi: 10.3389/fpsyg.2018.01123

3. Filimon F, Philiastides MG, Nelson JD, Kloosterman NA, Heekeren HR. How embodied is perceptual decision making? Evidence for separate processing of perceptual and motor decisions. Journal of Neuroscience. 2013 Jan 30;33(5):2121–36. doi: 10.1523/JNEUROSCI.2334-12.2013

4. Aczel B, Szollosi A, Palfi B, Szaszi B, Kieslich PJ. Is action execution part of the decision-making process? An investigation of the embodied choice hypothesis. Journal of Experimental Psychology Learning Memory and Cognition. 2018;44(6):918–926. doi: 10.1037/xlm0000484

5. Marcos E, Cos I, Girard B, Verschure PF. Motor cost influences perceptual decisions. PLoS One. 2015 Dec 16;10(12):e0144841. doi: 10.1371/journal.pone.0144841

6. Hagura N, Haggard P, Diedrichsen J. Perceptual decisions are biased by the cost to act. Elife. 2017 Feb 21;6:e18422. doi: 10.7554/eLife.18422

7. de Lange FP, Fritsche M. Perceptual decision-making: picking the low-hanging fruit?. Trends in cognitive sciences. 2017 May 1;21(5):306–7. doi: 10.1016/j.tics.2017.03.006

8. Burk D, Ingram JN, Franklin DW, Shadlen MN, Wolpert DM. Motor effort alters changes of mind in sensorimotor decision making. PLoS One. 2014 Mar 20;9(3):e92681. doi: 10.1371/journal.pone.0092681

9. Harris CM, Wolpert DM. The main sequence of saccades optimizes speed-accuracy trade-off. Biological cybernetics. 2006 Jul 1;95(1):21–9. doi: 10.1007/s00422-006-0064-

10. Armel KC, Beaumel A, Rangel A. Biasing simple choices by manipulating relative visual attention. Judgment and Decision making. 2008;3(5):396–403.

11. Lim SL, O’Doherty JP, Rangel A. The decision value computations in the vmPFC and striatum use a relative value code that is guided by visual attention. Journal of Neuroscience. 2011 Sep 14;31(37):13214–23. doi: 10.1523/JNEUROSCI.1246-11.2011

12. Atalay AS, Bodur HO, Rasolofoarison D. Shining in the center: Central gaze cascade effect on product choice. Journal of Consumer Research. 2012 May 3;39(4):848–66. doi: 10.1086/665984

13. Bird GD, Lauwereyns J, Crawford MT. The role of eye movements in decision making and the prospect of exposure effects. Vision Research. 2012 May 1;60:16–21. doi: 10.1016/j.visres.2012.02.014

14. Kunar MA, Watson DG, Tsetsos K, Chater N. The influence of attention on value integration. Attention, Perception, & Psychophysics. 2017 Aug 1;79(6):1615–27. doi: 10.3758/s13414-017-1340-7

15. Krajbich I, Armel C, Rangel A. Visual fixations and the computation and comparison of value in simple choice. Nature neuroscience. 2010 Oct;13(10):1292. doi: 10.1038/nn.2635

16. Krajbich I, Rangel A. Multialternative drift-diffusion model predicts the relationship between visual fixations and choice in value-based decisions. Proceedings of the National Academy of Sciences. 2011 Aug 16;108(33):13852–7. doi: 10.1073/pnas.1101328108

17. Tavares G, Perona P, Rangel A. The attentional drift diffusion model of simple perceptual decision-making. Frontiers in neuroscience. 2017 Aug 24;11:468. doi: 10.3389/fnins.2017.00468

18. Shimojo S, Simion C, Shimojo E, Scheier C. Gaze bias both reflects and influences preference. Nature neuroscience. 2003 Dec;6(12):1317. doi: 10.1038/nn1150

19. Krajbich I. Accounting for attention in sequential sampling models of decision making. Current opinion in psychology. 2018 Oct 13. doi: 10.1016/j.copsyc.2018.10.008

20. Gottlieb J. Understanding active sampling strategies: Empirical approaches and implications for attention and decision research. Cortex. 2018 May 1;102:150–60. doi: 10.1016/j.cortex.2017.08.019

21. Ludwig CJ, Evens DR. Information foraging for perceptual decisions. Journal of Experimental Psychology: Human Perception and Performance. 2017 Feb;43(2):245. doi: 10.1037/xhp0000299

22. Smith SM, Krajbich I. Gaze amplifies value in decision making. Psychological science. 2019 Jan;30(1):116–28. doi: 10.1177/0956797618810521

23. Hagura N, Diedrichsen J, Haggard P. Action cost biases the perceptual decision making, only when the cost is implicit. Translational and Computational Motor Control, 2013.

24. Wispinski NJ, Gallivan JP, Chapman CS. Models, movements, and minds: bridging the gap between decision making and action. Annals of the New York Academy of Sciences. 2018 Oct 1. doi: 10.1111/nyas.13973

25. Shadmehr R, Huang HJ, Ahmed AA. A representation of effort in decision-making and motor control. Current biology. 2016 Jul 25;26(14):1929–34. doi: 10.1016/j.cub.2016.05.065

26. Morel P, Ulbrich P, Gail A. What makes a reach movement effortful? Physical effort discounting supports common minimization principles in decision making and motor control. PLoS biology. 2017 Jun 6;15(6):e2001323. doi: 10.1371/journal.pbio.2001323

27. Orquin JL, Loose SM. Attention and choice: A review on eye movements in decision making. Acta psychologica. 2013 Sep 1;144(1):190–206. doi: 10.1016/j.actpsy.2013.06.003

28. Schulte-Mecklenbeck M, Johnson JG, Böckenholt U, Goldstein DG, Russo JE, Sullivan NJ, Willemsen MC. Process-tracing methods in decision making: On growing up in the 70s. Current Directions in Psychological Science. 2017 Oct;26(5):442–50. doi: 10.1177/0963721417708229

29. Heitz RP. The speed-accuracy tradeoff: history, physiology, methodology, and behavior. Frontiers in neuroscience. 2014 Jun 11;8:150. doi: 10.3389/fnins.2014.00150

30. Forstmann BU, Ratcliff R, Wagenmakers EJ. Sequential sampling models in cognitive neuroscience: Advantages, applications, and extensions. Annual review of psychology. 2016 Jan 4;67:641–66. doi: 10.1146/annurev-psych-122414-033645

31. Cassey TC, Evens DR, Bogacz R, Marshall JA, Ludwig CJ. Adaptive sampling of information in perceptual decision-making. PloS one. 2013 Nov 27;8(11):e78993. doi: 10.1371/journal.pone.0078993

32. Mullett TL, Stewart N. Implications of visual attention phenomena for models of preferential choice. Decision. 2016 Oct;3(4):231. doi: 10.1037/dec0000062

33. Onuma T, Penwannakul Y, Fuchimoto J, Sakai N. The effect of order of dwells on the first dwell gaze bias for eventually chosen items. PloS one. 2017 Jul 19;12(7):e0181641. doi: 10.1371/journal.pone.0181641

34. Dutilh G, Rieskamp J. Comparing perceptual and preferential decision making. Psychonomic bulletin & review. 2016 Jun 1;23(3):723–37. doi: 10.3758/s13423-015-0941-1

35. Heckerman D, Breese JS. Causal independence for probability assessment and inference using Bayesian networks. IEEE Transactions on Systems, Man, and Cybernetics-Part A: Systems and Humans. 1996 Nov;26(6):826–31. doi: 10.1109/3468.541341

36. Friedman N. Inferring cellular networks using probabilistic graphical models. Science. 2004 Feb 6;303(5659):799–805. doi: 10.1126/science.1094068

37. Yu J, Smith VA, Wang PP, Hartemink AJ, Jarvis ED. Advances to Bayesian network inference for generating causal networks from observational biological data. Bioinformatics. 2004 Jul 29;20(18):3594–603. doi: 10.1093/bioinformatics/bth448

