## Supplementary material for "Motor effort and adaptive sampling in perceptual decision-making": S1 Supporting Information. Mathematical details of the Bayesian Network modeling.

#### Mathematical Details of the Bayesian Network Modeling

#### Model (a)

##### -Control

The joint probability function for the network is:

$$\begin{aligned} &p(\text{right chosen}, \text{right last sampled}, \boldsymbol{DV}) \\ =&p(\text{right chosen}|\text{right last sampled}, \boldsymbol{DV})p(\text{right last sampled})p(\boldsymbol{DV}) \end{aligned} \tag{1}$$

We marginalize $\boldsymbol{DV}$:

$$\begin{aligned} &p(\text{right chosen}, \text{right last sampled}) \\ =&\int p(\text{right chosen}|\text{right last sampled}, \boldsymbol{DV})p(\text{right last sampled})p(\boldsymbol{DV})d\boldsymbol{DV} \end{aligned} \tag{2}$$

The conditional probability is:

$$\begin{aligned} &p(\text{right chosen}|\text{right last sampled}) \\ =&\int p(\text{right chosen}|\text{right last sampled}, \boldsymbol{DV})p(\boldsymbol{DV})d\boldsymbol{DV} \end{aligned} \tag{3}$$

##### -One-Switch

Let $\boldsymbol{SP}$ denote the start position in the trial. The joint probability function is:

$$\begin{aligned} &p(\text{right chosen}, \text{right last sampled}, \boldsymbol{DV}, \boldsymbol{SP}) \\ =&p(\text{right chosen}|\text{right last sampled}, \boldsymbol{DV})p(\text{right last sampled}|\boldsymbol{SP})p(\boldsymbol{DV})p(\boldsymbol{SP}) \end{aligned} \tag{4}$$

We marginalize $\boldsymbol{DV}$ and $\boldsymbol{SP}$:

$$
\begin{aligned}
&p(\text{right chosen}, \text{right last sampled}) \\
=&\iint p(\text{right chosen}|\text{right last sampled}, \boldsymbol{DV})p(\text{right last sampled}|\boldsymbol{SP})p(\boldsymbol{DV})p(\boldsymbol{SP})d\boldsymbol{DV}d\boldsymbol{SP} \\
=&\int p(\text{right chosen}|\text{right last sampled}, \boldsymbol{DV})p(\boldsymbol{DV})d\boldsymbol{DV}\int p(\text{right last sampled}|\boldsymbol{SP})p(\boldsymbol{SP})d\boldsymbol{SP} \\
=&\int p(\text{right chosen}|\text{right last sampled}, \boldsymbol{DV})p(\text{right last sampled})p(\boldsymbol{DV})d\boldsymbol{DV}
\end{aligned}
\tag{5}
$$

Therefore, the joint probability $p(\text{right chosen}, \text{right last sampled})$ is the same as in the Control condition, so is the conditional probability $p(\text{right chosen}|\text{right last sampled})$.

We assume that

$$
p(\text{right chosen}|\text{right last sampled}, \boldsymbol{DV}) = f(\boldsymbol{DV}) = \frac{1}{1+e^{-(a+b\boldsymbol{DV})}} \tag{6}
$$

where $a > 0.5$ and $b > 0$.

$$
\begin{aligned}
&p(\text{right chosen}|\text{right last sampled}) \\
=&\int p(\text{right chosen}|\text{right last sampled}, \boldsymbol{DV})p(\boldsymbol{DV})d\boldsymbol{DV} \\
=&\frac{1}{2}\int f(\boldsymbol{DV})p(\boldsymbol{DV})d\boldsymbol{DV} + \frac{1}{2}\int f(-\boldsymbol{DV})p(-\boldsymbol{DV})d\boldsymbol{DV} \\
=&\frac{1}{2}\int (f(\boldsymbol{DV}) + f(-\boldsymbol{DV}))p(\boldsymbol{DV})d\boldsymbol{DV}
\end{aligned}
\tag{7}
$$

We calculate the following derivative:

$$
\begin{aligned}
&\frac{d(f(\boldsymbol{DV}) + f(-\boldsymbol{DV}))}{d\boldsymbol{DV}} \\
=&bf(\boldsymbol{DV})(1-f(\boldsymbol{DV})) - bf(-\boldsymbol{DV})(1-f(-\boldsymbol{DV})) \\
=&b(\frac{1}{1+e^{-a-b\boldsymbol{DV}}}\frac{1}{1+e^{a+b\boldsymbol{DV}}} - \frac{1}{1+e^{-a+b\boldsymbol{DV}}}\frac{1}{1+e^{a-b\boldsymbol{DV}}}) \\
=&b(\frac{e^{a+b\boldsymbol{DV}}}{(1+e^{a+b\boldsymbol{DV}})^2} - \frac{e^{a+b\boldsymbol{DV}}}{(e^a+e^{b\boldsymbol{DV}})^2}) \\
=&-be^{a+b\boldsymbol{DV}}\frac{(e^{2a}-1)(e^{2b\boldsymbol{DV}}-1)}{(1+e^{a+b\boldsymbol{DV}})^2(e^a+e^{b\boldsymbol{DV}})^2}
\end{aligned}
\tag{8}
$$

When $\boldsymbol{DV} = 0$, $\frac{d(f(\boldsymbol{DV}) + f(-\boldsymbol{DV}))}{d\boldsymbol{DV}} = 0$.
When $\boldsymbol{DV} < 0$, $\frac{d(f(\boldsymbol{DV}) + f(-\boldsymbol{DV}))}{d\boldsymbol{DV}} > 0$; when $\boldsymbol{DV} > 0$, $\frac{d(f(\boldsymbol{DV}) + f(-\boldsymbol{DV}))}{d\boldsymbol{DV}} < 0$.

Therefore, $f(\boldsymbol{DV}) + f(-\boldsymbol{DV}))$ has a global maximum at $\boldsymbol{DV} = 0$ and decreases as $|\boldsymbol{DV}|$ increases, thus $\int (f(\boldsymbol{DV}) + f(-\boldsymbol{DV}))p(\boldsymbol{DV})d\boldsymbol{DV}$ increases as the variance of $p(\boldsymbol{DV})$ decreases.

**-Right-Biased**

The joint probability function with right-biased choice buttons is:

$$\begin{aligned} &p(\text{right chosen}, \text{right last sampled}, \boldsymbol{DV}, \text{Right-Biased}) \\ =&p(\text{right chosen}|\text{right last sampled}, \boldsymbol{DV})p(\text{right last sampled}|\text{Right-Biased})p(\boldsymbol{DV})p(\text{Right-Biased}) \end{aligned} \tag{9}$$

We marginalize $\boldsymbol{DV}$:

$$\begin{aligned} &p(\text{right chosen}, \text{right last sampled}, \text{Right-Biased}) \\ =&\int p(\text{right chosen}|\text{right last sampled}, \boldsymbol{DV})p(\text{right last sampled}|\text{Right-Biased})p(\boldsymbol{DV})p(\text{Right-Biased})d\boldsymbol{DV} \end{aligned} \tag{10}$$

The conditional probability is:

$$\begin{aligned} &p(\text{right chosen}|\text{right last sampled}, \text{Right-Biased}) \\ =&\int p(\text{right chosen}|\text{right last sampled}, \boldsymbol{DV})p(\boldsymbol{DV})d\boldsymbol{DV} \\ =&p(\text{right chosen}|\text{right last sampled}) \end{aligned} \tag{11}$$

### Model (b)

**-Control**

The joint probability function for the network is:

$$\begin{aligned} &p(\text{right chosen}, \text{right last sampled}, \boldsymbol{DV}) \\ =&p(\text{right chosen}|\boldsymbol{DV})p(\text{right last sampled}|\boldsymbol{DV})p(\boldsymbol{DV}) \end{aligned} \tag{12}$$

We marginalize $\boldsymbol{DV}$:

$$
\begin{aligned}
&p(\text{right chosen}, \text{right last sampled}) \\
&= \int p(\text{right chosen}|\boldsymbol{DV})p(\text{right last sampled}|\boldsymbol{DV})p(\boldsymbol{DV})d\boldsymbol{DV}
\end{aligned} \tag{13}
$$

The conditional probability is:

$$
\begin{aligned}
&p(\text{right chosen}|\text{right last sampled}) \\
&= \int p(\text{right chosen}|\boldsymbol{DV})\frac{p(\text{right last sampled}|\boldsymbol{DV})}{p(\text{right last sampled})}p(\boldsymbol{DV})d\boldsymbol{DV}
\end{aligned} \tag{14}
$$

**-One-Switch**

The joint probability function is:

$$
\begin{aligned}
&p(\text{right chosen}, \text{right last sampled}, \boldsymbol{DV}, \boldsymbol{SP}) \\
=&p(\text{right chosen}|\boldsymbol{DV})p(\text{right last sampled}|\boldsymbol{SP})p(\boldsymbol{DV})p(\boldsymbol{SP})
\end{aligned} \tag{15}
$$

We marginalize $\boldsymbol{DV}$ and $\boldsymbol{SP}$:

$$
\begin{aligned}
&p(\text{right chosen}, \text{right last sampled}) \\
&= \iint p(\text{right chosen}|\boldsymbol{DV})p(\text{right last sampled}|\boldsymbol{SP})p(\boldsymbol{DV})p(\boldsymbol{SP})d\boldsymbol{DV}d\boldsymbol{SP} \\
&= \int p(\text{right chosen}|\boldsymbol{DV})p(\text{right last sampled})p(\boldsymbol{DV})d\boldsymbol{DV}
\end{aligned} \tag{16}
$$

The conditional probability is:

$$
p(\text{right chosen}|\text{right last sampled}) = \int p(\text{right chosen}|\boldsymbol{DV})p(\boldsymbol{DV})d\boldsymbol{DV} \tag{17}
$$

**-Right-Biased**

The joint probability function with right-biased choice buttons is:

$$
\begin{aligned}
&p(\text{right chosen}, \text{right last sampled}, \boldsymbol{DV}, \text{Right-Biased}) \\
=&p(\text{right chosen}|\boldsymbol{DV})p(\text{right last sampled}|\boldsymbol{DV}, \text{Right-Biased})p(\boldsymbol{DV})p(\text{Right-Biased})
\end{aligned} \tag{18}
$$

We marginalize $\boldsymbol{DV}$:

$$\begin{aligned} & p(\text{right chosen}, \text{right last sampled}, \text{Right-Biased}) \\ = & \int p(\text{right chosen}|\boldsymbol{DV}) p(\text{right last sampled}|\boldsymbol{DV}, \text{Right-Biased}) p(\boldsymbol{DV}) p(\text{Right-Biased}) d\boldsymbol{DV} \end{aligned} \tag{19}$$

The conditional probability is:

$$\begin{aligned} & p(\text{right chosen}|\text{right last sampled}, \text{Right-Biased}) \\ = & \int p(\text{right chosen}|\boldsymbol{DV}) p(\text{right last sampled}|\boldsymbol{DV}, \text{Right-Biased}) \frac{p(\text{Right-Biased})}{p(\text{right last sampled}, \text{Right-Biased})} p(\boldsymbol{DV}) d\boldsymbol{DV} \\ = & \int p(\text{right chosen}|\boldsymbol{DV}) \frac{p(\text{right last sampled}|\boldsymbol{DV}, \text{Right-Biased})}{p(\text{right last sampled}|\text{Right-Biased})} p(\boldsymbol{DV}) d\boldsymbol{DV} \end{aligned} \tag{20}$$

We assume that the tendency to sample the right item last due to the right-biased choice buttons is independent of $\boldsymbol{DV}$, thus:

$$\begin{cases} p(\text{right last sampled}|\boldsymbol{DV}, \text{Right-Biased}) = p(\text{right last sampled}|\boldsymbol{DV}) + p_0 \\ p(\text{right last sampled}|\text{Right-Biased}) = p(\text{right last sampled}) + p_0 \end{cases} \tag{21}$$

where $p_0$ is the additional likelihood to sample the right item last.

Therefore,

$$\frac{p(\text{right last sampled}|\boldsymbol{DV}, \text{Right-Biased})}{p(\text{right last sampled}|\text{Right-Biased})} = \frac{p(\text{right last sampled}|\boldsymbol{DV}) + p_0}{p(\text{right last sampled}) + p_0} \tag{22}$$

When $\boldsymbol{DV} > 0$, $p(\text{right last sampled}|\boldsymbol{DV}) > p(\text{right last sampled})$; when $\boldsymbol{DV} < 0$, $p(\text{right last sampled}|\boldsymbol{DV}) < p(\text{right last sampled})$, thus:

$$\begin{cases} \frac{p(\text{right last sampled}|\boldsymbol{DV}) + p_0}{p(\text{right last sampled}) + p_0} < \frac{p(\text{right last sampled}|\boldsymbol{DV})}{p(\text{right last sampled})}, \boldsymbol{DV} > 0 \\ \frac{p(\text{right last sampled}|\boldsymbol{DV}) + p_0}{p(\text{right last sampled}) + p_0} > \frac{p(\text{right last sampled}|\boldsymbol{DV})}{p(\text{right last sampled})}, \boldsymbol{DV} < 0 \end{cases} \tag{23}$$

so $p(\text{right chosen}|\text{right last sampled}, \text{Right-Biased}) < p(\text{right chosen}|\text{right last sampled})$.

### Model (c)

**-Control**

The joint probability function for the network is:

$$\begin{aligned}&p(\text{right chosen}, \text{right last sampled}, \boldsymbol{DV})\\=&p(\text{right chosen}|\text{right last sampled}, \boldsymbol{DV})p(\text{right last sampled}|\boldsymbol{DV})p(\boldsymbol{DV})\end{aligned} \tag{24}$$

We marginalize $\boldsymbol{DV}$:

$$\begin{aligned}&p(\text{right chosen}, \text{right last sampled})\\=&\int p(\text{right chosen}|\text{right last sampled}, \boldsymbol{DV})p(\text{right last sampled}|\boldsymbol{DV})p(\boldsymbol{DV})d\boldsymbol{DV}\end{aligned} \tag{25}$$

The conditional probability is:

$$\begin{aligned}&p(\text{right chosen}|\text{right last sampled})\\=&\int p(\text{right chosen}|\text{right last sampled}, \boldsymbol{DV})\frac{p(\text{right last sampled}|\boldsymbol{DV})}{p(\text{right last sampled})}p(\boldsymbol{DV})d\boldsymbol{DV}\end{aligned} \tag{26}$$

**-One-Switch**

The joint probability function is:

$$\begin{aligned}&p(\text{right chosen}, \text{right last sampled}, \boldsymbol{DV}, \boldsymbol{SP})\\=&p(\text{right chosen}|\text{right last sampled}, \boldsymbol{DV})p(\text{right last sampled}|\boldsymbol{SP})p(\boldsymbol{DV})p(\boldsymbol{SP})\end{aligned} \tag{27}$$

We marginalize $\boldsymbol{DV}$ and $\boldsymbol{SP}$:

$$\begin{aligned}&p(\text{right chosen}, \text{right last sampled})\\=&\iint p(\text{right chosen}|\text{right last sampled}, \boldsymbol{DV})p(\text{right last sampled}|\boldsymbol{SP})p(\boldsymbol{DV})p(\boldsymbol{SP})d\boldsymbol{DV}d\boldsymbol{SP}\\=&\int p(\text{right chosen}|\text{right last sampled}, \boldsymbol{DV})p(\text{right last sampled})p(\boldsymbol{DV})d\boldsymbol{DV}\end{aligned} \tag{28}$$

The conditional probability is:

$$p(\text{right chosen}|\text{right last sampled}) = \int p(\text{right chosen}|\text{right last sampled}, \boldsymbol{DV})p(\boldsymbol{DV})d\boldsymbol{DV} \tag{29}$$

**-Right-Biased**

The joint probability function with right-biased choice buttons is:

$$\begin{aligned}&p(\text{right chosen}, \text{right last sampled}, \boldsymbol{DV}, \text{Right-Biased})\\ =&p(\text{right chosen}|\text{right last sampled}, \boldsymbol{DV})p(\text{right last sampled}|\boldsymbol{DV}, \text{Right-Biased})p(\boldsymbol{DV})p(\text{Right-Biased})\end{aligned} \tag{30}$$

We marginalize $\boldsymbol{DV}$:

$$\begin{aligned}&p(\text{right chosen}, \text{right last sampled}, \text{Right-Biased})\\ =&\int p(\text{right chosen}|\text{right last sampled}, \boldsymbol{DV})p(\text{right last sampled}|\boldsymbol{DV}, \text{Right-Biased})p(\boldsymbol{DV})p(\text{Right-Biased})d\boldsymbol{DV}\end{aligned} \tag{31}$$

The conditional probability is:

$$\begin{aligned}&p(\text{right chosen}|\text{right last sampled}, \text{Right-Biased})\\ =&\int p(\text{right chosen}|\text{right last sampled}, \boldsymbol{DV})\frac{p(\text{right last sampled}|\boldsymbol{DV}, \text{Right-Biased})}{p(\text{right last sampled}|\text{Right-Biased})}p(\boldsymbol{DV})d\boldsymbol{DV}\\ <&p(\text{right chosen}|\text{right last sampled})\end{aligned} \tag{32}$$
